# Dehydrozaluzanin C- derivative protects septic mice by alleviating over-activated inflammatory response and promoting the phagocytosis of macrophages

**DOI:** 10.1101/2023.11.01.565192

**Authors:** Ying-xiang Zou, Tian-nan Xiang, Li-rong Xu, Huan Zhang, Yu-he Ma, Lu Zhang, Chun-xian Zhou, Xiao Wu, Qi-lin Huang, Biao Lei, Jing-wen Mu, Xiang-yang Qin, Xin Jiang, Yue-juan Zheng

## Abstract

Host-directed therapy (HDT) is a new adjuvant strategy that interfere with host cell factors that are required by a pathogen for replication or persistence. In this study, we assessed the effect of dehydrozaluzanin C-derivative (DHZD), a modified compound from dehydrozaluzanin C (DHZC), as a potential HDT agent for severe infection. LPS-induced septic mouse model and Carbapenem resistant *Klebsiella pneumoniae* (CRKP) infection mouse model was used for testing *in vivo*. RAW264.7 cells, mouse primary macrophages, and DCs were used for *in vitro* experiments. Dexamethasone (DXM) was used as a positive control agent. DHZD ameliorated tissue damage (lung, kidney, and liver) and excessive inflammatory response induced by LPS or CRKP infection in mice. Also, DHZD improved the hypothermic symptoms of acute peritonitis induced by CRKP, inhibited heat-killed CRKP (HK-CRKP)-induced inflammatory response in macrophages, and upregulated the proportions of phagocytic cell types in lungs. *In vitro* data suggested that DHZD decreases LPS-stimulated expression of IL-6, TNF-α and MCP-1 via PI3K/Akt/p70S6K signaling pathway in macrophages. Interestingly, the combined treatment group of DXM and DHZD had a higher survival rate and lower level of IL-6 than those of the DXM-treated group; the combination of DHZD and DXM played a synergistic role in decreasing IL-6 secretion in sera. Moreover, the phagocytic receptor CD36 was increased by DHZD in macrophages, which was accompanied by increased bacterial phagocytosis in a clathrin- and actin-dependent manner. This data suggests that DHZD may be a potential drug candidate for treating bacterial infections.

## Introduction

Sepsis is a life-threatening organ dysfunction caused by a dysregulated response to infection (1, 2). Approximately 15 to 30 million cases of sepsis are reported annually (3, 4). Although age-standardized sepsis incidence has dropped by 37.0% and mortality decreased by 52.8% from 1990 to 2017 (5), antimicrobial resistance still challenges the treatment of sepsis. With the increasing use of carbapenems in hospitals, CRKP has become a major threat to public health. CRKP accounts for about 64% of carbapenem-resistant enterobacterial infections in China (6, 7). Although a number of studies have reported that the combined therapy with multiple active agents may be effective for CRKP strains, the rate of clinical failure remains high (8–10). Thus, identifying new effective treatment strategies for CRKP is crucial.

The current clinical treatment for sepsis caused by bacterial infection mainly relies on antibiotic therapy to control the infection and the use of glucocorticoids to inhibit the excessive inflammatory response. Glucocorticoids, such as DXM, are effective but also accelerate specific immune cells’ apoptosis. Common side effects of long-term glucocorticoid use include osteoporosis (11) and opportunistic infections due to its immunosuppressive effects (12). The development of new antimicrobial drugs, especially those that target G^-^ bacteria, lags behind the emergence of antibiotic resistance, which has become a major obstacle in the clinic.

As a major component of the cell wall of G^-^ bacteria, LPS commonly known as endotoxin, is one of the most important pathogen-associated molecular patterns, which is sensed by our immune system via the Toll-like receptor 4 (TLR4) (13). This ligation activates adaptor molecules such as myeloid differentiation protein 88 (MyD88), interleukin-1 (IL-1) receptor-associated kinases (IRAKs), tumor necrosis factor (TNF) receptor-associated factor 6 (TRAF6), transforming growth factor (TGF)-activated kinase 1 (TAK1), and subsequent MAPKs and nuclear factor κB (NF-κB). Yet, excessive secretion of inflammatory cytokines (such as IL-6, TNF-α and IL-Iβ) can lead to “cytokine storm”, commonly known as systemic inflammatory response syndrome (SIRS). The pathologic progress of sepsis is usually accompanied by SIRS. The uncontrolled inflammatory response may lead to organ dysfunction, multiple organ failure, and, in more severe cases, even death (14).

Besides the inflammatory immune damage, the poor clearance of pathogens is usually associated with poor prognosis during bacterial infection. Monocytes/macrophages and dendritic cells (DCs) are the main phagocytes that phagocytize bacteria and present their antigens to T cells (15). Neutrophil is also an important cell type of phagocytes. Phagocytosis includes three main stages: receptor-mediated, membrane fusion, and actin-driven (16). During this process, phagocytic receptors, including C-type lectin receptors (CLRs), mannose receptors, scavenger receptors (SR-A1, MARCO and CD36), complement receptors (CR3 and CR4) and Fc receptors, have a crucial role (16–18).

HDT is a new adjuvant strategy often used to treat severe infections that can overcome AMR. It is not mediated by killing pathogens directly; instead, it promotes the immune clearing of pathogens through phagocytes and/or reduces tissue damage caused by excessive inflammatory response (19).

DHZC, a sesquiterpenoid from *Magnolia officinalis*, exhibits strong anti-inflammatory activity, but its drug toxicity is relatively high. In this study, we assessed the effect of DHZD, a modified compound from DHZC, as a potential HDT agent for severe infection with anti-inflammatory and immunomodulatory effects.

## Materials and Methods

### Animals and reagents

Six-week-old wild-type female C57BL/6J mice (18 ± 2 g) were obtained from Joint Ventures Sipper BK Experimental Animal Co. (Shanghai, China). All mice had free access to food and water during all the experiments and were housed in a 12/12 h light/dark cycle. The experimental procedures were approved by the Experimental Animal Ethical Committee, Shanghai University of Traditional Chinese Medicine (Shanghai, China) (PZSHUTCM220307018).

DHZD was modified from DHZC isolated from *Magnolia officinalis*. LPS (0111: B4), DMSO, dexamethasone and chlorpromazine hydrochloride (CPZ) were purchased from Sigma (St. Louis, MO, USA). Cell counting kit 8 (CCK-8) were obtained from Meilunbio (Dalian, China). Enzyme-linked immunosorbent assay (ELISA) kits for mouse IL-6, TNF-α, MCP-1, IL-10, IL-1β, IFN-β and granulocyte-macrophage colony stimulating factor (mGM-CSF) were purchased from

R&D Systems (Minneapolis, MN, USA). Mouse recombinant cytokine IL-4 were from Peprotech (USA). The detection kit of nitric oxide (NO) was obtained from Beyotime Biotechnology (Shanghai, China). FITC anti-mouse CD11c, APC anti-mouse CD40, APC/Fire™ 750 anti-mouse CD80, PerCP/Cyanine5.5 anti-mouse CD86, Brilliant Violet 421™ anti-mouse I-A/I-E, FITC anti-mouse CD45, Brilliant Violet 650 Ly6G, Brilliant Violet 421 Ly6C and PE/Cyanine7 F4/80 were purchased from Biolegend (USA). Dexamethasone sodium phosphate injection was from Sinopharm (China). SYBR RT-PCR Kit was purchased from Takara (Dalian, China). PHrodo-labelled *E. coli* was purchased from ThermoFisher (USA). Cytochalasin D was obtained from Abcam (USA). Filipin III was from MCE (USA). Levofloxacin (LVX) and meropenem (MEM) were obtained from the National Institute for Food and Drug Control of China (Beijing, China) CRKP (HS11286) was kindly provided by Dr. Yi-jian Chen from Huashan Hospital affiliated to Fudan University.

### Cell culture

Mouse macrophage-like cell line RAW264.7 were purchased from ATCC (Manassas, VA, USA) and cultured as previously described (20).

Thioglycolate-elicited mouse primary peritoneal macrophages were isolated from female C57BL/6J mice at 6-8 weeks of age and cultured in DMEM (GE, USA) with 10 % FBS (Biowest, France) and 1 % Penicillin-Streptomycin, as previously described (21).

Bone marrow-derived macrophages (BMDMs) and bone marrow-derived dendritic cells (BMDCs) were both obtained from C57BL/6J mice (4 weeks old). The BMDMs were cultured in DMEM containing 10 ng/mL M-CSF (Peprotech, USA) and 10 % FBS. The BMDCs were differentiated in RPMI 1640 (GE, USA) medium containing 10 % FBS, 50 ng/mL mGM-CSF (R&D Systems, USA) and 4 ng/mL mIL-4 (Peprotech, USA).

### Cell viability assay

RAW264.7 cells (1×10^5^ cells/well) were cultured in 96-well plates overnight and treated with different concentrations of DHZD (0-15 μM) in 12, 24, 48 and 72 h. The cytotoxicity of DHZD was evaluated by CCK-8 counting kit according to the manufacturer’s instructions (Meilunbio, Dalian, China).

### Preparation of HK-CRKP

Monoclonal colonies were picked from the CRKP bacterial culture plates and cultured in LB medium for 12 h at 37°C on a constant temperature shaker. After heating 1 mL of the bacterial solution in a 65°C water bath for 45 min, the solution was centrifuged (4400 rpm×15 min), resuspended in sterile PBS, centrifuged again (4400 rpm×15 min), and resuspended in DMEM for further use (MOI=10).

### Enzyme-linked immunosorbent assay

RAW264.7 (2×10^5^ cells/well) were plated as indicated and administrated with DHZD and/or LPS (100 ng/mL) for 6 h and 18 h. The levels of cytokines (IL-6, TNF-α, MCP-1, IFN-β, IL-1β and IL-10) in the cell culture supernatants were detected by ELISA (R&D Systems, USA).

Mouse primary macrophages were plated (3.5 × 10^5^ cells/500 µL) in 24-well plates at the day before stimulation. Cell culture supernatants were collected at 6 h and 18 h after co-stimulation of different concentrations of DHZD and LPS (100 ng/mL), and the concentrations of IL-6, TNF-α, MCP-1, IL-1β, IFN-β and IL-10 were measured by ELISA.

BMDCs were re-seeded on day 5 of differentiation at 3.5 × 10^5^ cells/500 μL in 24-well plates and stimulated by DHZD (0-15 μM) with or without LPS (100 ng/mL) for 6 h or 18 h. The secretion of IL-6, TNF-α and IL-12p70 in cell culture supernatants were detected by ELISA.

Mouse primary macrophages were plated (3.5 × 10^5^ cells/500 µL) in 24-well plates one day before and were stimulated by different concentrations of DHZD and HK-CRKP (MOI=10). After 12 h or 24 h, cell culture supernatants were collected and the concentrations of IL-6, TNF-α, MCP-1, IL-1β, IL-10 and MIP-2 were measured by ELISA.

BMDMs were re-seeded on day 5 of differentiation (3.5 × 10^5^ cells/well) in 24-well plates and then stimulated by LPS (100 ng/mL) and DHZD for 6 h. The expression of cytokines (IL-6, TNF-α and IL-10) was detected by ELISA.

### Assessment of NO level

Mouse primary macrophages were plated (3.5 × 10^5^ cells/500 µL) in 24-well plates at the day before stimulation. Cell culture supernatants were collected at 6 h and 18 h after co-stimulation of different concentrations of DHZD and LPS (100 ng/mL). The secretion of NO was detected by corresponding NO detection kit (S0021) which was purchased from Beyotime Biotechnology (Shanghai, China).

### Detection of MHC II and co-stimulatory molecules on BMDCs

Bone marrow stem cells were derived and differentiated into BMDCs. After 5 days, BMDCs were seeded in 12-well plates overnight. LPS (100 ng/mL) and DHZD (15 μM) were added into culture medium of BMDCs for another 24 h. Then BMDCs were harvested and labeled with FITC anti-mouse CD11c (Biolegend, 117305, USA), APC anti-mouse CD40 (Biolegend, 124611, USA), APC/Fire™ 750 anti-mouse CD80 (Biolegend, 104739, USA), PerCP/Cyanine5.5 anti-mouse CD86 (Biolegend, 105027, USA) and Brilliant Violet 421™ anti-mouse I-A/I-E (Biolegend, 107631, USA). And then cells were analyzed by Flow cytometry (Beckman, USA), and data were analyzed using FlowJo software.

### The establishment of septic mice by lethal dose of LPS challenge

Six groups were included (n=10): LPS group, LPS+DXM group (5 mg/kg), LPS+DHZD group (10 mg/kg), LPS+DHZD group (20 mg/kg), LPS+DHZD group (40 mg/kg) and LPS+MEM (5 mg/kg) +DHZD group (10 mg/kg). The septic mouse model was established by intraperitoneal injection of LPS (12.5 mg/kg, Sigma) as described (22, 23). At the same time, DHZD and DXM were injected intraperitoneally.

Survival status of different groups was recorded every 2-3 h in the following continuous 180 h (24).

### LPS-induced acute peritonitis

Acute peritonitis mouse model (n=9) was carried out by intraperitoneal injection of LPS (10 mg/kg). DXM (5 mg/kg) and/or DHZD (10 or 40 mg/kg) were administered at the same time. After 12 h, all the mice in different groups were sacrificed. Meanwhile, blood samples were collected and allowed to clot for 3 h followed by centrifugation at 3500 rpm for 25 min (4 [) to prepare serum samples. Serum levels of cytokines (IL-6, TNF-α, MCP-1 and IL-10) were measured by corresponding ELISA kits (R&D Systems, USA) according to the manufacturer’s instructions.

Total RNA was extracted from lung tissues and reversely transcribed to cDNA. Real-time PCR was performed with SYBR RT-PCR kit (Takara, Dalian, China). The mRNA levels were normalized against β-Actin and presented as 2^-ΔΔCt^.

The lung tissues were snipped and digested with collagenase (Sigma Aldrich) and DNase I (Sigma Aldrich) in gentleMACS^TM^ (Miltenyi, Germany) at 80 rpm (37 °C) for 30 min. After digestion, the samples were filtered through a 40 mm filter using a syringe piston, and centrifuged at 1000 rpm (4 °C) for 5 min. RBC lysis buffer (BD Biosciences, USA) was added to cell pellets at room temperature for 5 min, and then PBS (8 mL) was added and centrifuged at 1400 rpm (4 °C) for 5 min. Cells were counted and then incubated at 4 °C for 25 min with FITC anti-mouse CD45 (Biolegend, 103107, USA), Brilliant Violet 650 Ly6G (Biolegend, 127641, USA),

Brilliant Violet 421 Ly6C (Biolegend, 128031, USA) and PE/Cyanine7 F4/80 (Biolegend, 123113, USA) antibodies. Samples were run on CytoFlex LX (Beckman, USA). Data were analyzed using FlowJo software.

Lung, liver and kidney tissues were collected to be fixed with 4% formaldehyde and paraffin-embedded. All of the above tissues were sliced and stained with hematoxylin and eosin (H&E). The histopathological changes were observed using a PANNORAMIC MIDI II microscope (3DHistech, Hungary). Pulmonary edema, alveolar infiltration, and pulmonary vasculitis were scored on a 12-point pathological scale (25).

### RNA extraction and real-time PCR

RAW264.7 cells were seeded (2×10^5^/well) in 24-well plates and incubated overnight. Cells were treated with heat killed-Methicillin resistant *Staphylococcus aureus* (HK-MRSA) (MOI=10) separately along with DHZD (15 μM) for 12 and 18 h. The cells were harvested for total RNA extraction and reversely transcribed to cDNA for further qPCR detection.

Real-time PCR was performed with SYBR RT-PCR kit (Takara, Dalian, China). The mRNA levels were normalized against β-Actin and presented as 2^-ΔΔCt^. The primer sequences for qRT-PCR amplification were as the following:

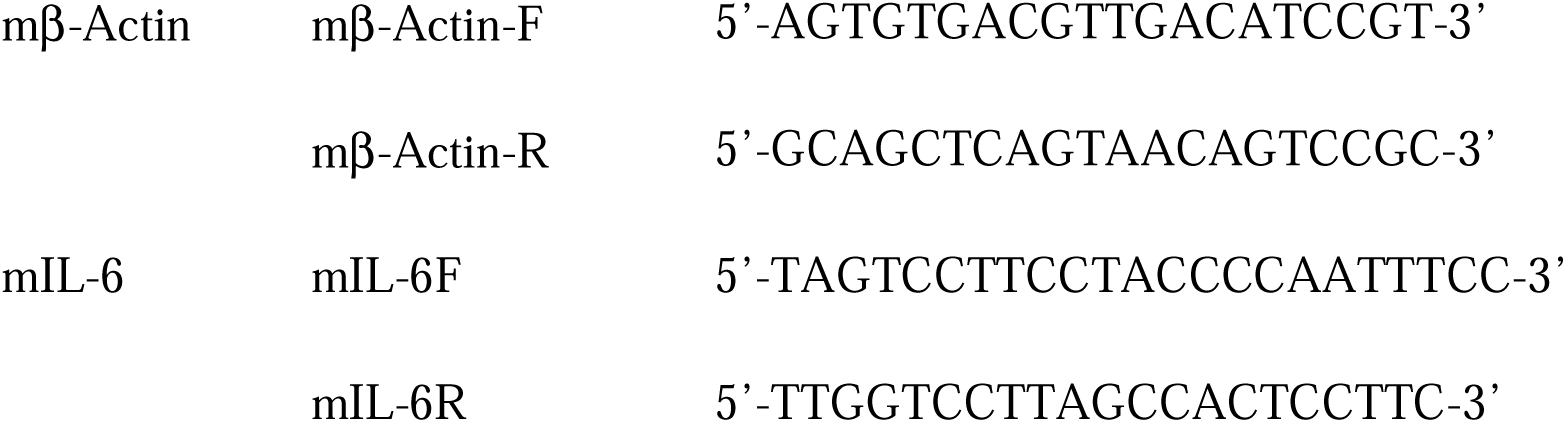
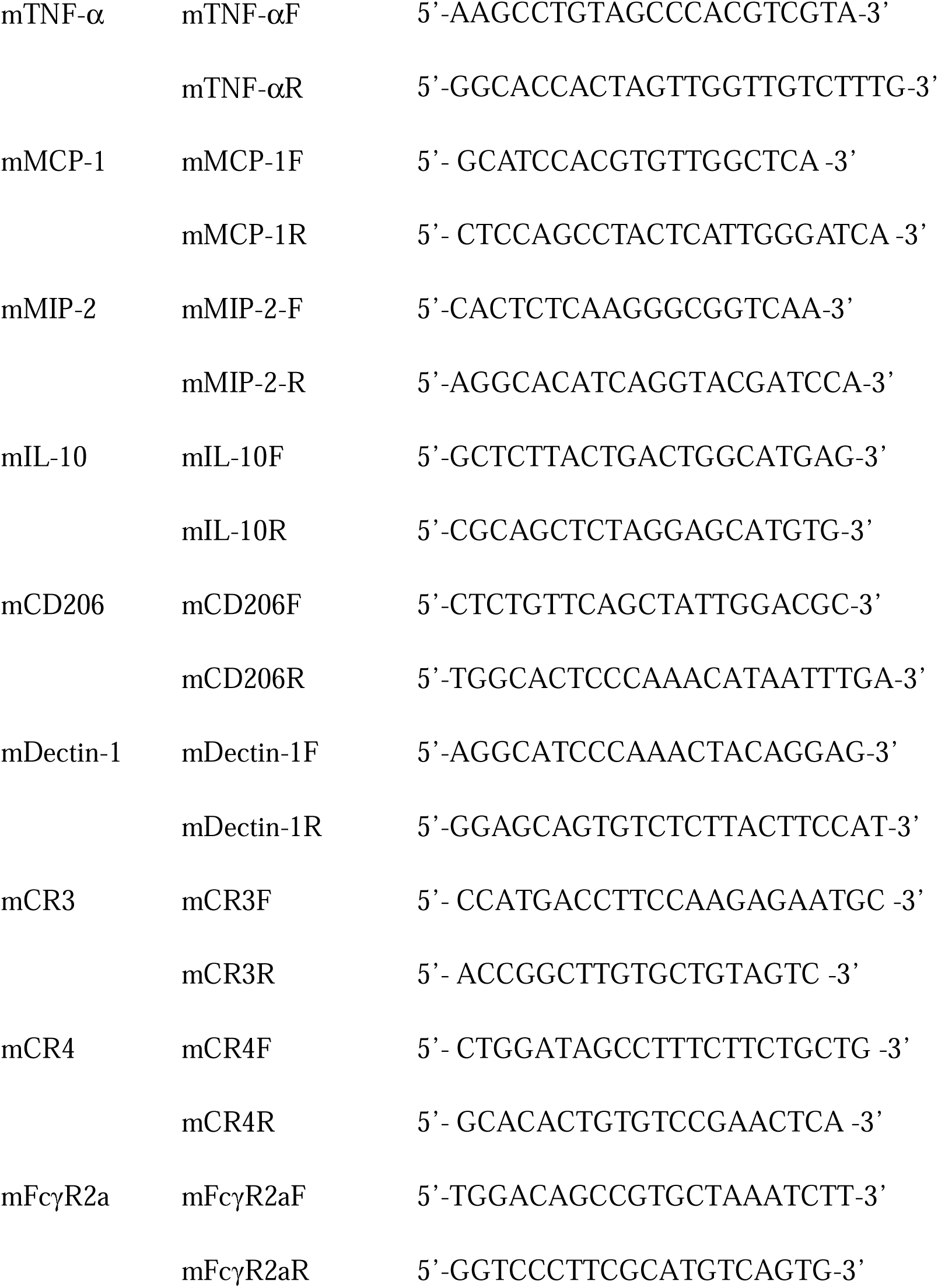

### Detection of the phagocytosis of macrophages

The phagocytosis of pHrodo labelled *E. coli* (ThermoFisher, California, USA) was detected in RAW264.7 cells by Flow cytometry. RAW264.7 cells were plated at 1×10^5^ cells/100 µL in 96-well plates and incubated overnight. Then cells were primed with LPS (100 ng/mL) along with DHZD (15 µM). After 24 h, macrophages were incubated with pHrodo-labelled *E. coli* (10 μL/well) protected from light for 1 h. Cells were washed with sterile PBS for 3 times to get rid of the bacteria outside of the cells. Then the phagocytosis of pHrodo-labelled *E. coli* was quantified by FACS. RAW264.7 cells were pretreated by 3 µM of Filipin III (the inhibitor of caveolin, MCE, USA), 10 µM of chlorpromazine hydrochloride (the inhibitor of clathrin, Sigma, USA) or 10 µM of cytochalasin D (the inhibitor of actin, Abcam, USA) for 30 min, and then pHrodo-labelled *E. coli* were added to incubation system for another 1 h. The phagocytosis of *E. coli* was investigated by BD Accuri^TM^ C6 Flow cytometry (BD, USA), and data were analyzed by FlowJo software.

### Western blot

The phosphorylation of important signaling proteins from RAW264.7 cells and their corresponding total proteins were determined by Western blot as previously described (26). RAW264.7 cells (1 × 10^6^ cells/well) were seeded in 6-well plates overnight and stimulated with LPS (100 ng/mL) with or without DHZD (15 μM) for a certain period of time. All the antibodies were bought from Cell Signaling Technology

(Beverly, MA, USA).

### CRKP-induced septic mouse model

The septic mouse model was established by intraperitoneal injection of CRKP (6×10^7^ CFU/mouse) in 0.4 mL of PBS. The experiments were carried out in the following groups: CRKP group, CRKP+MEM (5 mg/kg) group, CRKP+LVX (75 mg/kg) group, CRKP+DHZD group (20 mg/kg), CRKP+DHZD group (40 mg/kg) and the combined treatment CRKP+DHZD (20 mg/kg) +MEM (5 mg/kg) group. All medicines were administered simultaneously. The body temperature of mice was recorded by infrared body thermometer (HJKIR, Wuhan) before and 9 hours after injection of CRKP.

After 12 h, mice were sacrificed, and lungs and livers were collected to be fixed with 4% formaldehyde and embedded in paraffin. All of the above tissues were sliced and stained with H&E. The histopathological changes were observed using a PANNORAMIC MIDI II microscope (3DHistech, Hungary). Pulmonary edema, alveolar infiltration, and pulmonary vasculitis were scored on a 12-point pathological scale (25).

### Statistical analysis

Statistical analysis was performed using GraphPad Prism 9. Data were presented as means ± standard deviation (SD). Student’s *t*-test analyses was used to evaluate the comparisons between two groups. Comparisons multiple groups were evaluated by one-way ANOVA followed by Tukey’s post hoc comparisons, respectively. *P* values were considered as *, *P*<0.05; **, *P*<0.01; ***, *P*<0.001; ****, *P*<0.0001.

## Results

### DHZD inhibits the secretion of LPS-induced cytokines from RAW264.7 cells

The structure of DHZD is shown in Figure 1A. DHZD was synthesized according to the previously reported method (27). Spectroscopic data were in accordance with reported values (Supplementary Fig.1) (27).

**Figure 1.**
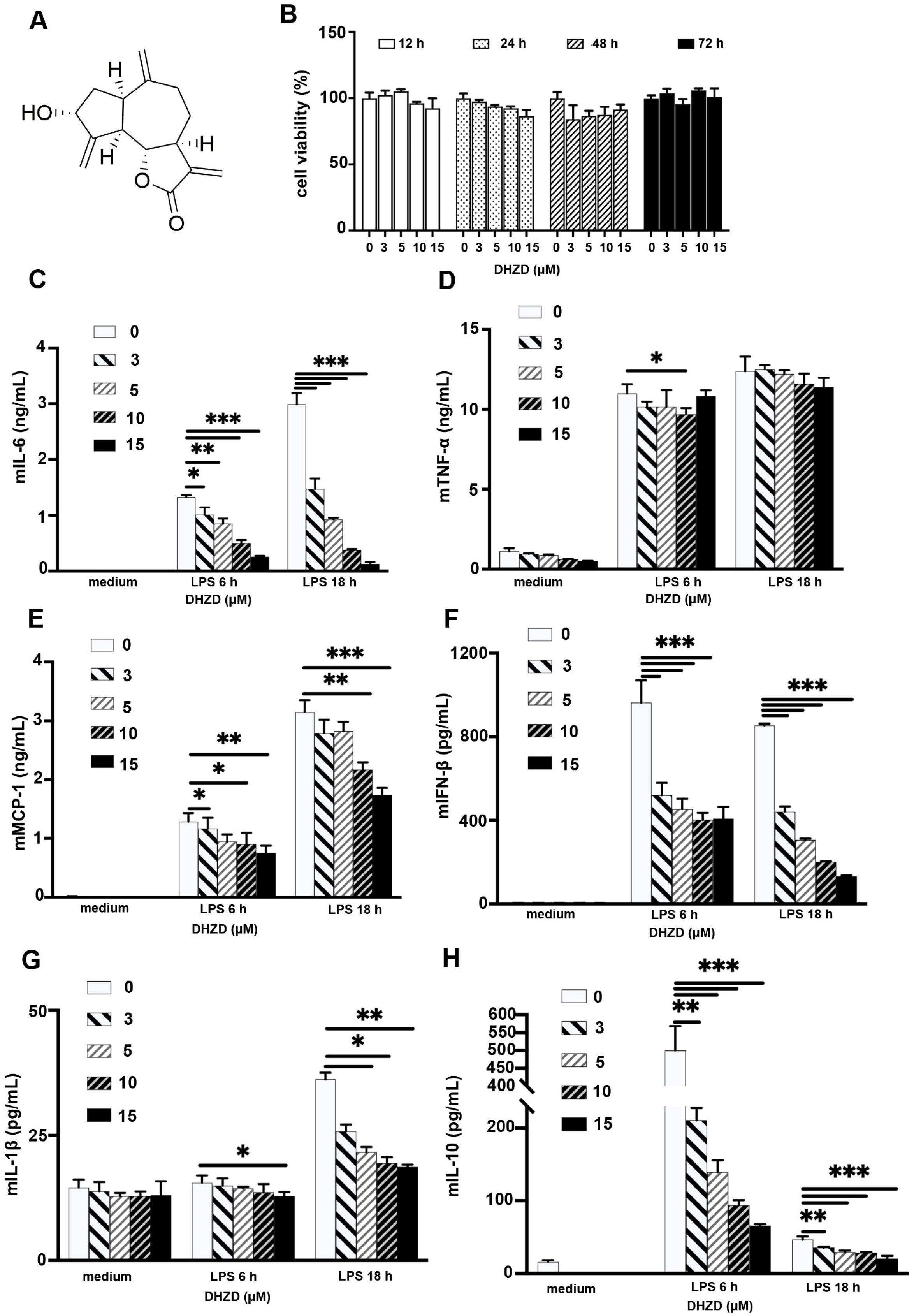
DHZD inhibits the secretion of LPS-induced cytokines from RAW264.7 cells. Chemical structure of DHZD (A). DHZD had no cytotoxicity on RAW264.7 cells (B). DHZD decreased the secretion of IL-6 (C), TNF-α (D), MCP-1 (E), IFN-β (F), IL-1β (G) and IL-10 (H) in the cell culture supernatants induced by LPS. Values are means of three replicate determinations (n=3) ± SD. *, *P*<0.05, **, *P* <0.01, and ***, *P* <0.001.

CCK-8 kit was used to determine whether DHZD affected cell viability. DHZD (0-15 μM) did not affect the proliferation of RAW264.7 cells within 72 h, but inhibited LPS-induced secretion of IL-6, TNF-α, MCP-1, IFN-β, and IL-10 in a dose-dependent manner (Fig. 1B-H).

### DHZD decreases the secretion of LPS-induced expression of IL-6, TNF-**α**, MCP-1, IL-1**β**, IFN-**β** and IL-10 in mouse primary macrophages

Next, we assessed the effect of DHZD on the secretion of LPS-induced expression of inflammatory mediators in mouse primary macrophages and BMDMs. Compared to macrophage-like cell lines, primary macrophages mimic the physiological state of cells *in vivo* and are expected to maintain their *in vivo* functions under optimal conditions for a short period (28). DHZD inhibited the secretion of LPS-induced pro-inflammatory mediators IL-6, TNF-α, IL-1β, IFN-β, NO, MCP-1 and anti-inflammatory cytokine IL-10 in mouse primary macrophages (Figure 2A-G) and decreased the secretion of IL-6, TNF-α and IL-10 in BMDMs stimulated with LPS (Supplementary Fig. 2A-C).

**Figure 2.**
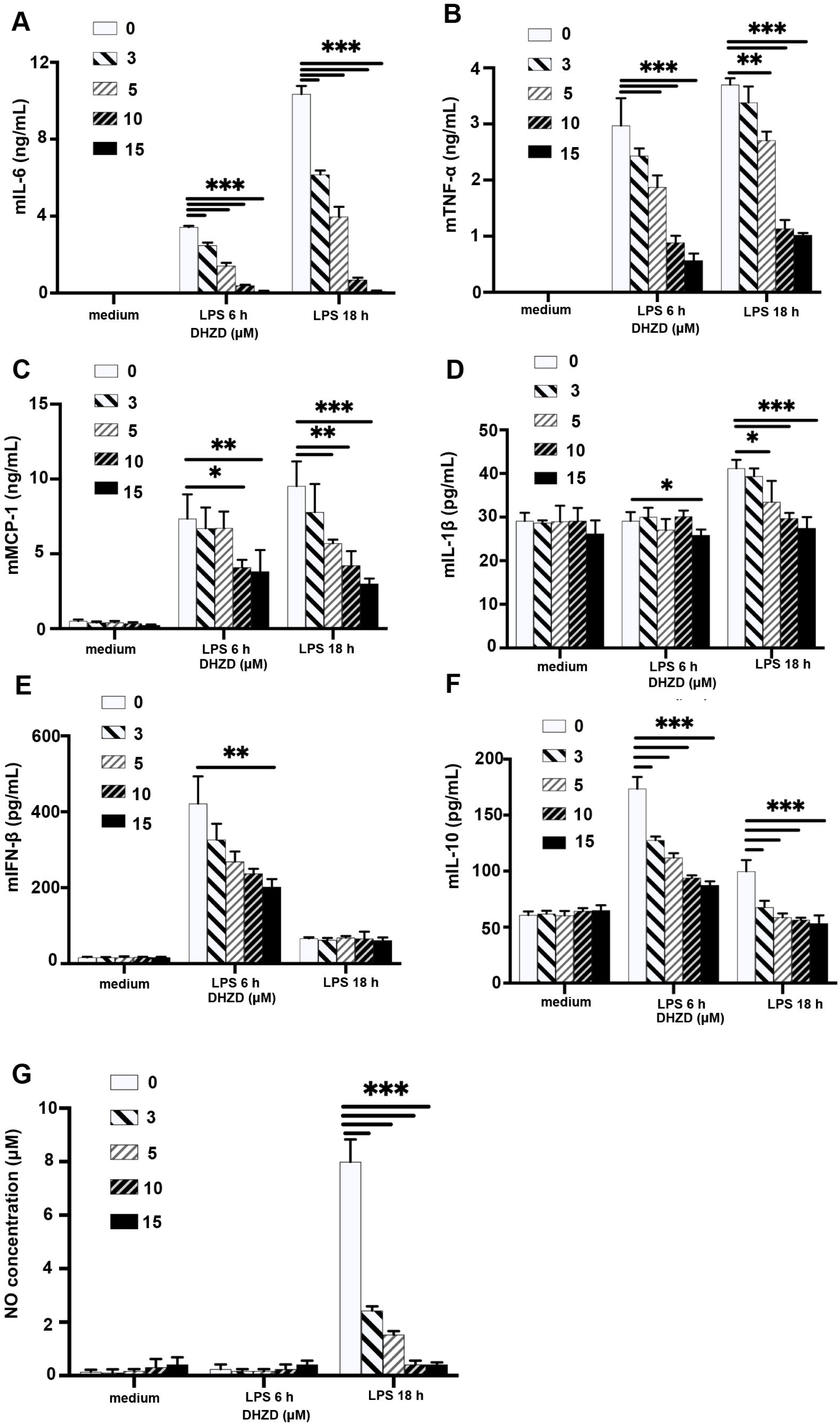
DHZD decreases the secretion of LPS-induced expression of IL-6, TNF-α, MCP-1, IL-1β, IFN-β and IL-10 in mouse primary macrophages. DHZD decreased LPS-induced secretion of IL-6 (A), TNF-α (B), MCP-1 (C), IL-1β (D), IFN-β (E), IL-10 (F) and NO (G). Data are means of three replicate determinations ± SD (n=3). *, *P* <0.05, **, *P* <0.01, and ***, *P* <0.001.

### DHZD decreases LPS-induced secretion of inflammatory cytokines in BMDCs, and downregulated the expression of MHC II and co-stimulatory molecules on the surface of BMDCs

DCs are the most important and powerful antigen-presenting cells that activate naive T cells, bridging innate and adaptive immunity. DCs participate in the phagocytosis and digestion of bacteria and present their antigens to evoke T-cell immunity (29). During the transition of DCs from immature to mature status, co-stimulatory molecules are upregulated, downregulating the phagocytosis and promoting the antigen-presenting ability. In order to investigate whether DHZD has a regulatory effect on the antigen presentation function of DCs, thereby affecting the activation of T cells, flow cytometry was used to detect the expression of costimulatory molecules (CD80, CD86, and CD40) and MHC Class II molecule (Iab) in cells treated with or without LPS. Briefly, DHZD inhibited the expression of these molecules compared to control cells (Figure 3D-G).

**Figure 3.**
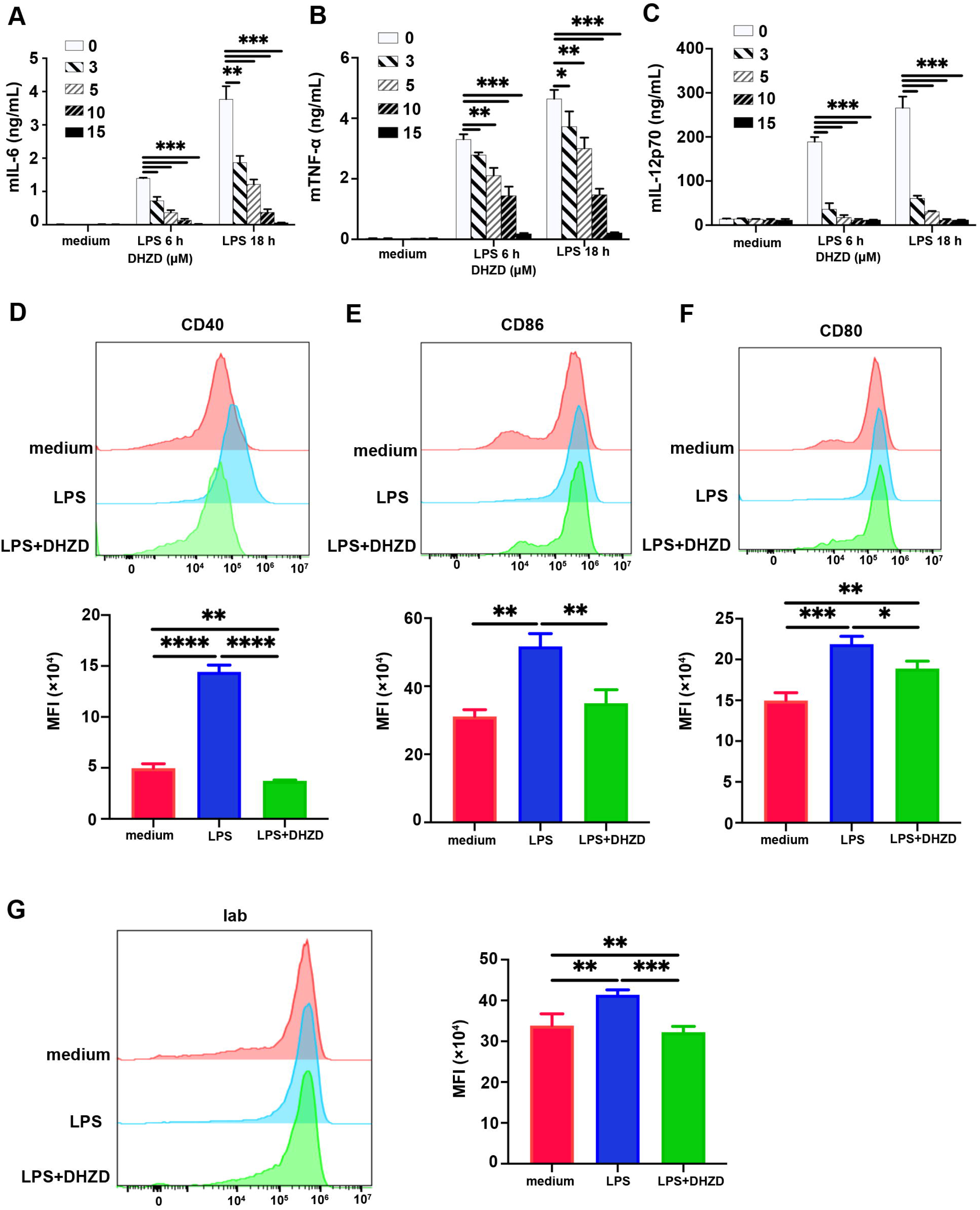
DHZD decreases LPS-induced secretion of inflammatory cytokines in BMDCs, and downregulated the expression of MHC II and co-stimulatory molecules on the surface of BMDCs. DHZD down-regulated the production of IL-6 (A), TNF-α (B) and IL-12p70 (C) in cell culture supernatants induced by LPS. The expression of co-stimulatory molecules CD40 (D), CD86 (E), CD80 (F) and antigen-presenting molecule MHC II (Iab) (G) was quantified by FACS and the MFI (mean fluorescence intensity) of each sample was analyzed using FlowJo. Data were shown as meanL±LSD of three separated experiments. *, *P*<0.05, **, *P* <0.01, ***, *P* <0.001, and ****, *P* <0.0001.

DCs secret inflammatory cytokines (IL-6, TNF-α, IL-12 and a variety of chemokines), contributing to the formation of cytokine storm together with activated T cells during the pathological progress of sepsis (30). This study found that DHZD reduces the secretion of IL-6, TNF-α, and IL-12p70 (Figure 3A-C) DCs stimulated with LPS.

### DHZD protects septic mice by inhibiting the over-activated inflammatory response

To determine the protective role of DHZD against LPS-induced (12.5 mg/kg) septic shock, a survival analysis was carried out. DXM, a glucocorticoid used in the clinical treatment of bacterial infection, was used as a positive control drug.

All mice in the model group died within 40 h. The mice in the septic model group had low body temperature, shivering, standing hair and decreased activity. In the DXM treatment group, 60 % of mice survived (10 % of mice in the low-dose DHZD treatment group and 30 % in the medium- or high-dose DHZD treatment groups). All mice survived in the low-dose DHZD and DXM treatment groups (Figure 4A). Compared with the model group, the mice in the DXM group, the high-dose DHZD-treated group, and the DXM+DHZD combination group exhibited faster recovery of body temperature, smoother hair, and were more active. The combined treatment group (DXM+DHZD) showed the best protective effect in terms of clinical manifestation.

**Figure 4.**
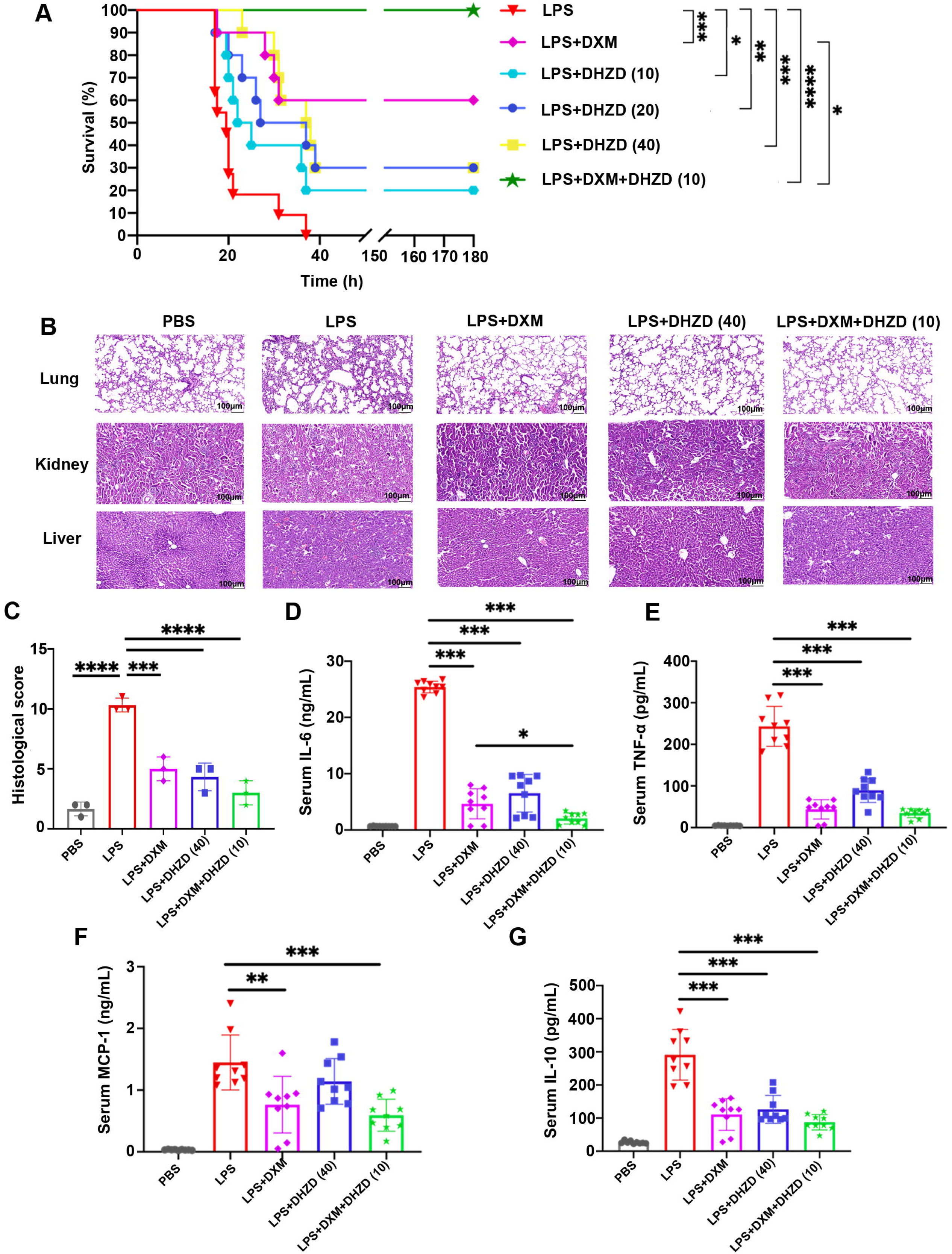
DHZD protects septic mice by inhibiting the over-activated inflammatory response. Survival (A). Data were analyzed using Log-Rank test and survival curve was generated by GraphPad. H&E staining of lung, kidney and liver (100×) (B). Scale bar 100 μm, and the histological core of lung (C). The serum levels of IL-6 (D), TNF-α (E), MCP-1 (F) and IL-10 (G) were measured by ELISA. Data were shown as mean ± SD of nine mice per group. *, *P* <0.05, **, *P* <0.01, ***, *P* <0.001, and ****, *P* <0.0001.

The lung, liver and kidney tissue structures in the control group were intact. On the contrary, obvious infiltration of inflammatory cells, a large amount of fluid exudation in the alveoli, and lung interstitial thickening were seen in mice injected with LPS. Also, the liver tissue showed obvious inflammatory cell infiltration, swelling and necrosis of hepatocytes, fragmentation and dissolution of some nuclei, and destruction of the hepatic plate structure, while renal tissue showed obvious inflammatory cell infiltration, degeneration and necrosis of renal tubular epithelial cells, and narrowing of renal cysts. DHZD improved pulmonary edema, alveolar infiltration, and lung vasculitis caused by LPS. DXM improved lung pathological damage (Figure 4B-C). The destruction of lung, liver and kidney tissue structure was improved after DXM, DHZD or DXM+DHZD treatment. There was no significant difference among different treated groups (DXM vs DHZD, *p*=0.8856, DXM vs DXM+DHZD, *p*=0.1168, DHZD vs DXM+DHZD, *p*=0.4112) (Figure 4B).

Multiple organ failure caused by a “cytokine storm” during infection is the leading cause of death in septic patients (31). In LPS-challenged mice, DHZD and DXM decreased the secretion of IL-6, TNF-α, MCP-1 and IL-10. Compared with DXM treatment, the combination treatment of DHZD+DXM could significantly inhibit the secretion of these cytokines (Figure 4D-G).

Sepsis induced by bacterial infection can cause systemic organ dysfunction, and lungs are often the most easily affected important organs. Therefore, sepsis is often accompanied by acute lung injury (ALI) (32) or acute respiratory distress syndrome (ARDS) (33), which is one of the most important prognostic factors of sepsis death. In this study, we detected the expression of lung inflammatory factors by qPCR. DHZD and DHZD+DXM could significantly inhibit the production of proinflammatory cytokines IL-6, chemokines MCP-1 and MIP-2 in the lungs of mice (Supplementary Fig. 3A-C).

### DHZD increases the numbers of lung neutrophils, monocytes, and macrophages induced by LPS

In most cases, locally recruited natural immune cells (such as neutrophils and monocytes/macrophages) can rapidly phagocyte bacteria, control the bacterial load in tissues, and reduce the occurrence of severe infection. However, when severe infection occurs, many immune cells participate in the secretion of inflammatory cytokines, forming cytokine storms, which can lead to organ damage, sepsis and even death. Thus, in the early stage of infection, rapid phagocytosis and elimination of bacteria, and subsequently control of the immune response of the body within reasonable limitation, are the key aspects for treatment (34, 35).

In this study, the infiltration of neutrophils and monocytes/macrophages in the lungs was measured by flow cytometry. As shown in Figure 5, 12 h after LPS injection, the ratio of neutrophils (Figure 5A-C) and monocytes/macrophages (Figure 5D-H) increased in the DXM-treated group, DHZD-treated group, and DHZD+DXM group compared to LPS challenged group, suggesting an increase in the number of innate immune cells with the ability to phagocyte bacteria.

**Figure 5.**
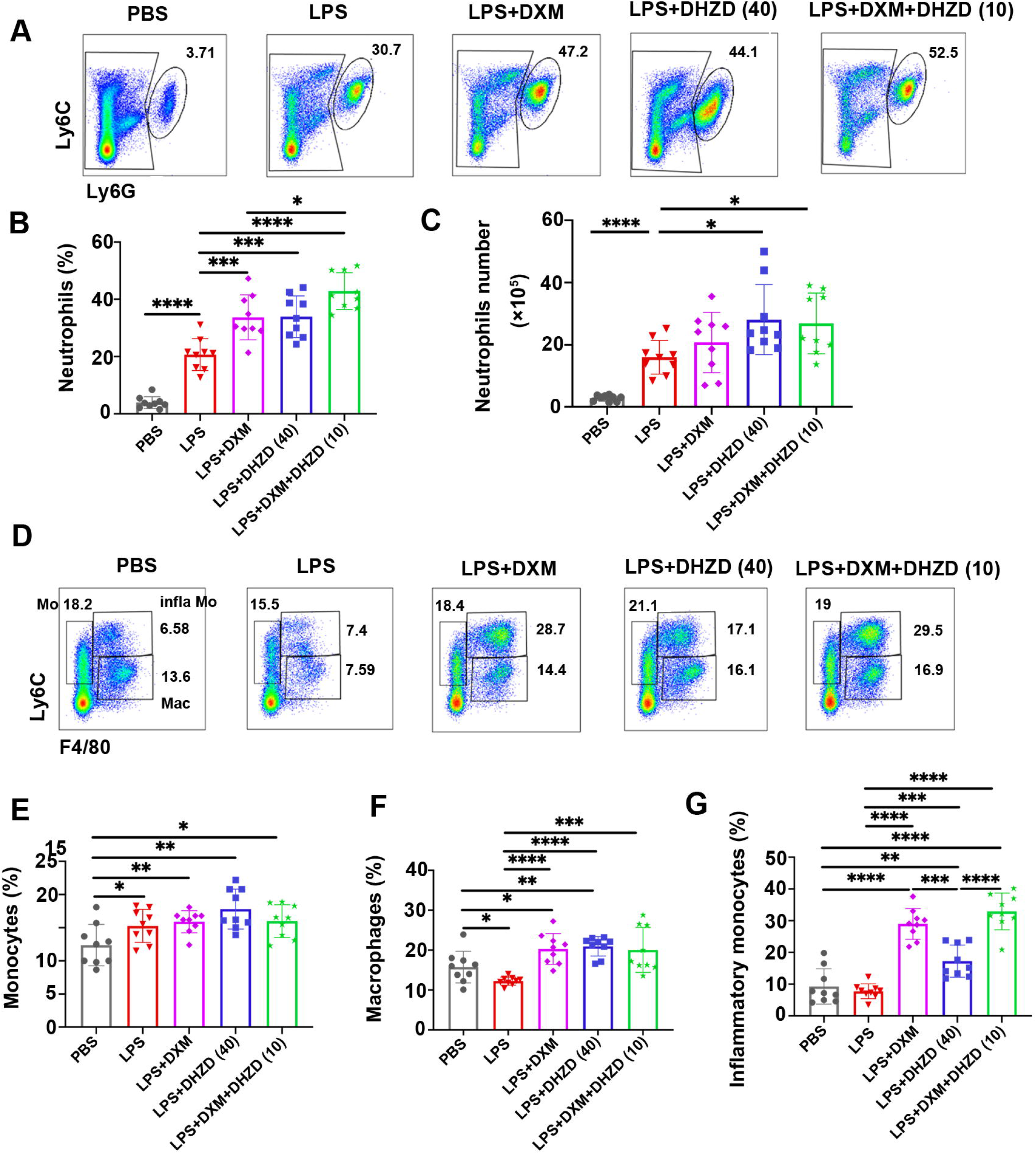
DHZD increases the numbers of lung neutrophils, monocytes, and macrophages induced by LPS. The proportions of lung neutrophils, monocytes and macrophages in lungs were detected. One representative scatter plot per group and the proportions of neutrophils (A-C). One representative scatter plot and the proportion of monocyte-macrophages (D-H). Data were shown as means ± SD for each group (nine mice in each group). *, *P*<0.05, **, *P* <0.01, ***, *P* <0.001, and ****, *P* < 0.0001.

DHZD may have rapid phagocytic role in the early stage of bacterial infection, reducing the bacterial load and the severity of infection in tissues, and thus exerting a potential protective role. From our results, DXM and DHZD have similar effects in increasing the proportions and numbers of neutrophils and monocytes. In addition, DXM also has a broad effect of immunosuppression, which might be accompanied by potential adverse effects (36).

### DHZD enhances the phagocytosis of pHrodo-labelled *E. coli* in macrophages in a clathrin-dependent and actin-polymerized manner via CD36

PHrodo-labeled *E. coli* is a model bacterial that emit fluorescence only in phagocytic cells because of the acidic environment (37, 38). To clarify the regulatory effect of DHZD on the phagocytotic ability, RAW264.7 cells were stimulated by LPS and treated with DHZD (15 µM) for 24 h. PHrodo-labeled *E. coli* was added and incubated for 1 h in dark. The amount of engulfed *E. coli* in the cytoplasm was measured by flow cytometry. DHZD significantly increased the phagocytosis of pHrodo-labeled *E. coli* in RAW264.7 (Figure 6A-C), suggesting that DHZD could promote the ability of bacterial phagocytosis of macrophages.

**Figure 6.**
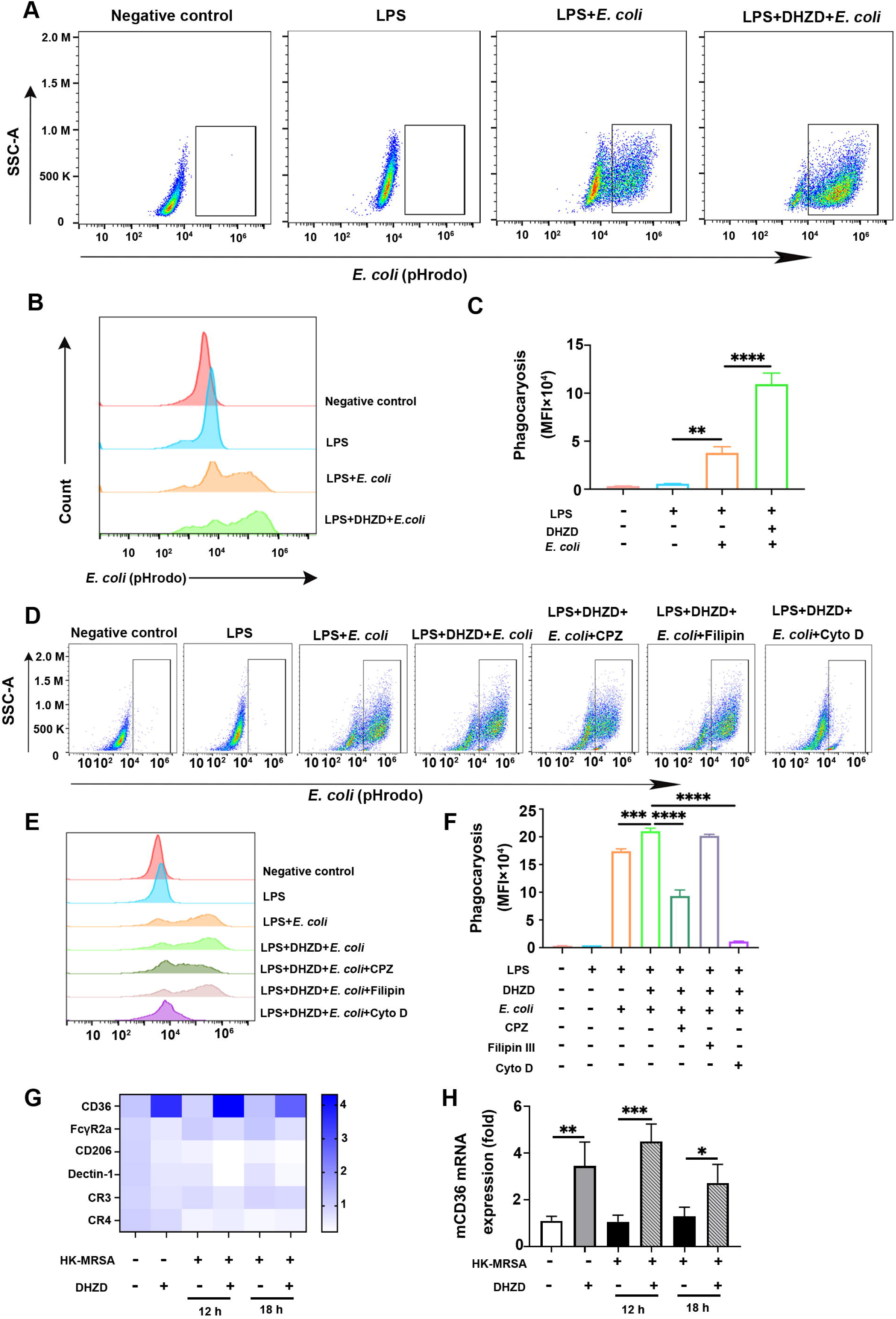
DHZD enhances the phagocytosis of pHrodo-labelled E. coli in macrophages in a clathrin-dependent and actin-polymerized manner via CD36. One representative scatter plot out of three independent experiments were shown (A). The mean fluorescence intensity (MFI) of pHrodo-labelled *E. coli* swallowed by RAW264.7 cells were analyzed (B, C). DHZD enhanced the phagocytosis in a clathrin-dependent and actin-polymerized manner (D-F). The expression of CD36 (G, H) were detected by PCR. Data were shown as mean ± SD from three independent experiments. *, *P* <0.05, **, *P* <0.01, ***, *P* <0.001, and ****, *P* <0.0001.

The initiation of phagocytosis depends on the recognition and binding of host cell receptors to the corresponding ligands on the surface of the pathogens. Innate immune cells such as macrophages, neutrophils and dendritic cells can phagocyte pathogens through caveolin-mediated endocytosis, clathrin-mediated endocytosis, pinocytosis and other ways (39, 40). All the above-mentioned phagocytotic mechanisms are actin-dependent. Filipin III, CPZ and cytochalasin D are inhibitors of caveolin, clathrin and actin, respectively. Our data show that Filipin III has no effect on DHZD-promoted phagocytosis of bacteria in RAW264.7 cells, while CPZ could partially inhibit DHZD-promoted phagocytosis. Cytochalasin D could prevent actin polymerization and completely prevent enhanced phagocytosis by DHZD (Figure 6D-F).

Phagocytic receptors usually mediate bacterial phagocytosis. The expression of phagocytic receptors CD36, FcγR2a, CD206, Dectin-1, CR3 and CR4 were detected by qRT-PCR on the surface of macrophages. From the expression screen, DHZD could promote the expression of CD36 (Figure 6G-H), at least partially accounting for the increased phagocytosis ability.

### DHZD inhibits LPS-stimulated activation of PI3K/Akt/p70S6K pathway

LPS, as a ligand of TLR4, can activate extracellular signal-regulated kinase 1/2 (ERK1/2), c-Jun N-terminal kinase (JNK), and p38 MAPK pathways, NF-κB pathway and phosphoinositide 3-kinase (PI3K)/Akt signaling pathway, leading to the production of cytokines (41). Inhibition of PI3K/Akt activation reduces LPS-induced release of NO, MCP-1, IL-6, and TNF-α (42). Our results demonstrated that DHZD can effectively reduce LPS-induced inflammatory response. DHZD inhibited LPS-induced phosphorylation of Akt at Ser473 and Thr308 and subsequent p70S6K at Thr389, but did not affect the activation of NF-κB, ERK and p38 MAPK signaling pathways (Figure 7A-E).

**Figure 7.**
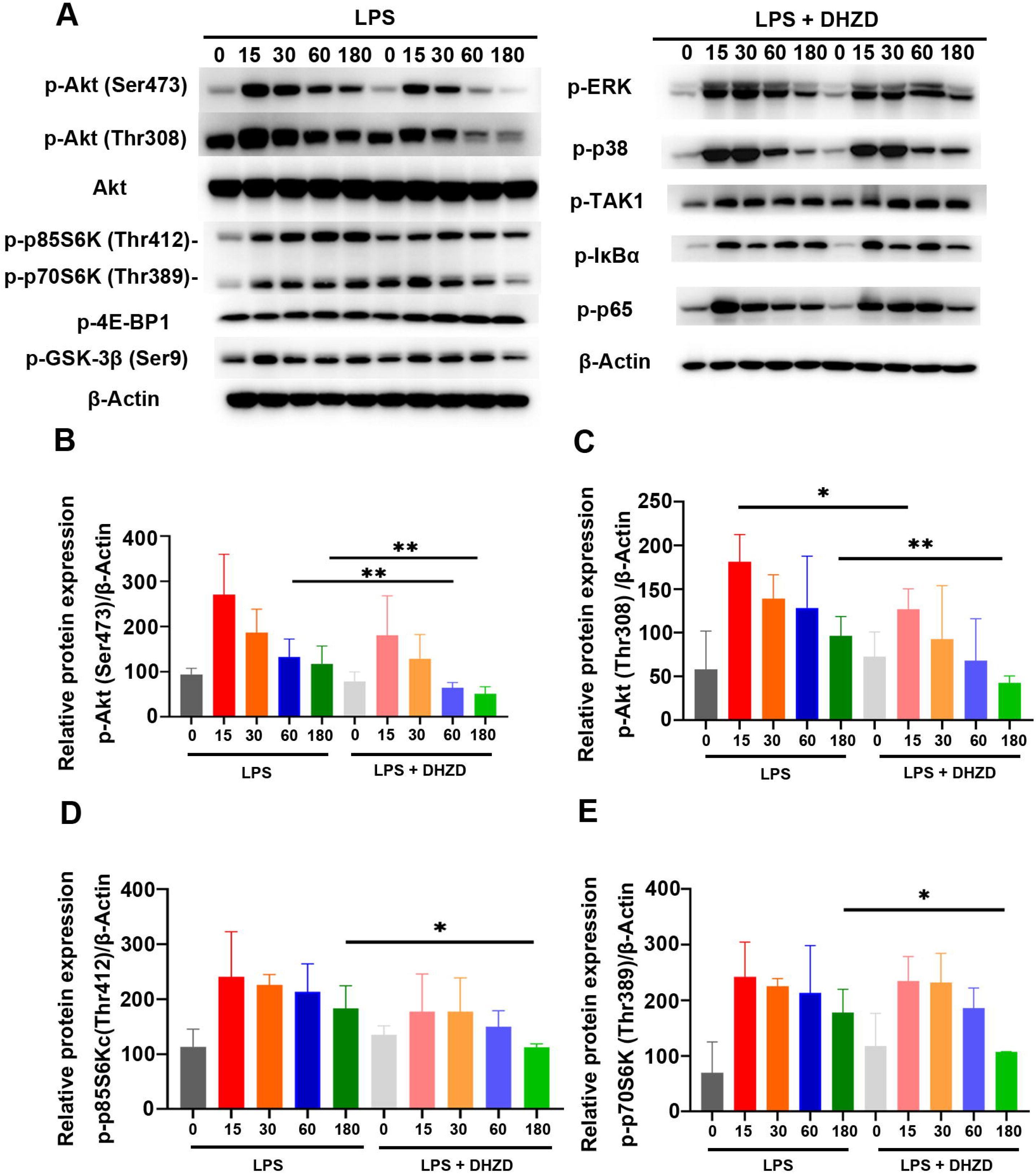
DHZD inhibits LPS-stimulated activation of PI3K/Akt/p70S6K pathway. Total protein of cells was detected by Western blot (A-E). Phospho-Akt (Ser 473), phospho-GSK-3β (Ser9), phospho-ERK, phospho-p38, phospho-IκBα (Ser32/36), phospho-p65 (Ser536) and β-Actin in RAW264.7 were verified by Western blot. Data were shown as mean ± SD from three independent experiments; *, *P* <0.05, **, *P* <0.01, ***, *P* <0.001, and ****, *P* <0.0001.

### DHZD reverses hypothermia, tissue damage, and over-activated inflammation caused by CRKP

LPS is the main component of the cell wall of G^-^ bacteria. CRKP is one of the most troublesome G^-^ bacteria causing severe infection and sepsis in the emergency department or intensive care unit (ICU). Fever is usually one of the most important symptoms of infection (43). There are variable thermoregulatory responses in sepsis, including fever and hypothermia (44). According to the analysis of clinical data, the mortality rate of septic patients with hypothermia is much higher than in those with fever (45).

In order to clarify the protective effects of DHZD on CRKP-infected mice, a septic mouse model was established by the intraperitoneal injection of CRKP, which is sensitive to LVX, used as a positive control antibiotic. Infrared thermal imaging thermometers were also used to monitor the body temperature of mice before and after the CRKP challenge. Compared with the model group, average body temperature was normalized by 1.3 [and 3.1 [in the DHZD (20) and DHZD (40) groups, respectively. The results suggested that DHZD had an impressive protective effect on the recovery of body temperature (Figure 8A, B).

**Figure 8.**
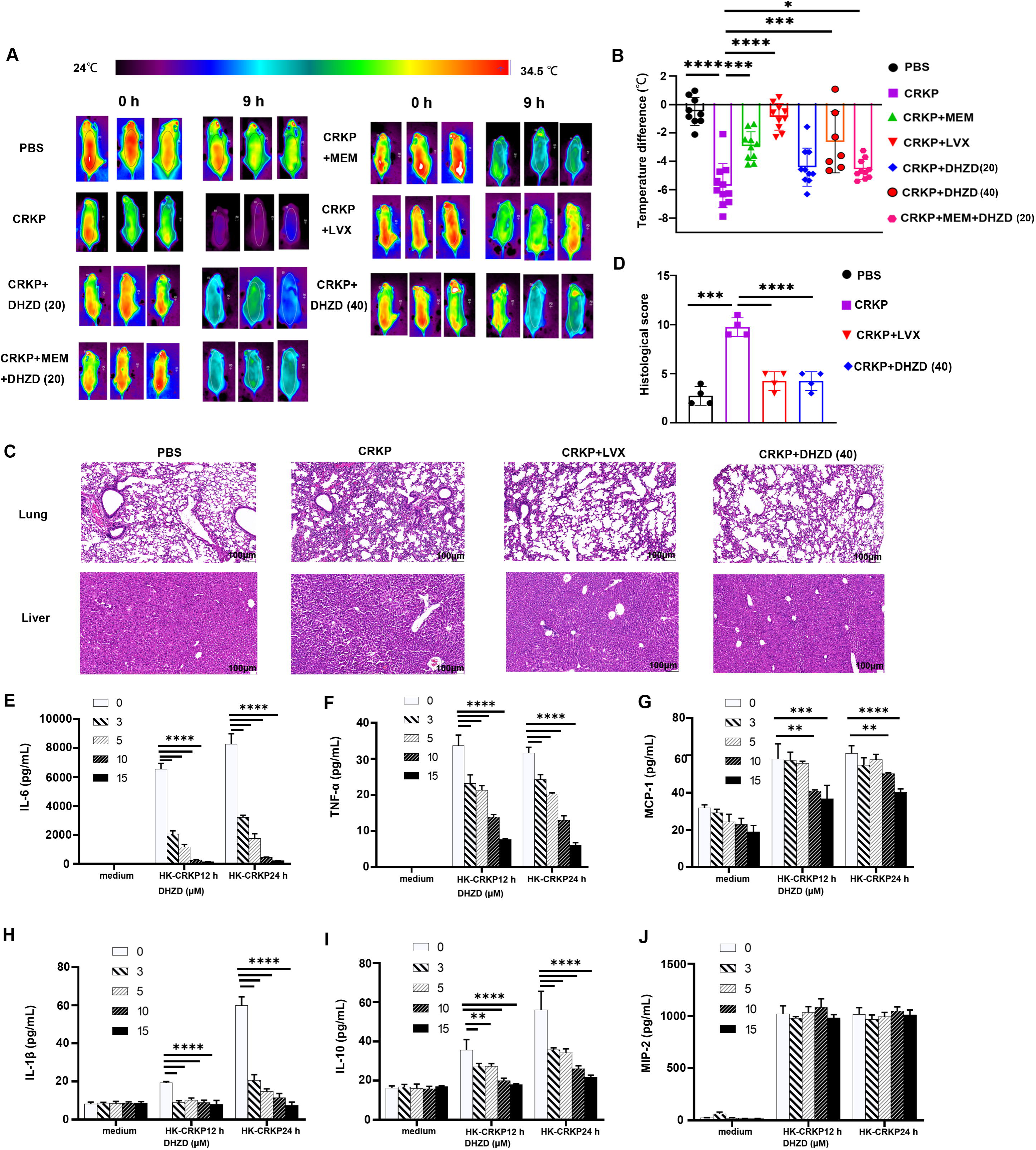
DHZD reverses hypothermia, tissue damage, and over-activated inflammation caused by CRKP. Temperature difference (A, B). H&E staining of lung and liver (200×) (C). Scale bar 100 μm, and the histological core of lung (D). DHZD dose-independently reduced the secretion of IL-6 (E), TNF-α (F), MCP-1 (G), IL-1β (H), IL-10 (I) and MIP-2 (J) induced by HK-CRKP. Data were shown as mean ± SD from three independent experiments. *, *P*<0.05, **, *P* <0.01, ***, *P* <0.001, and ****, *P* <0.0001.

Also, compared with the PBS group, the lung tissue of mice in the CRKP group showed obvious infiltration of inflammatory cells, a large amount of fluid exudation in the alveoli, and pulmonary interstitial thickening. Inflammatory cell infiltration, swelling and necrosis of hepatocytes, fragmentation and lysis of some nuclei, and destruction of hepatic plate structure were observed in the liver. DHZD ameliorated lung and liver tissue damage. A pathological score of lung tissue, as measured by pulmonary edema, alveolar infiltration, and pulmonary vasculitis, showed a protective effect of DHZD administration against pathological damage to the tissue (Figure 8C-D).

*In vitro* stimulation of primary peritoneal macrophages of mice with HK-CRKP showed that DHZD could reduce the secretion of IL-6, TNF-α, MCP-1, IL-1β and IL-10 in a dose-dependent manner but had no obvious influence on MIP-2 secretion (Figure 8E-J).

## Discussion

The World Health Organization has listed AMR among top 10 threats for global health. After entering the bloodstream, invading bacteria are recognized by pathogen-associated molecular patterns (PAMPs) via pattern recognition receptors (PRRs). In most cases, the innate immune response eliminated the invading bacteria.

Yet, when the bacteria prevail in the host, the immune response becomes unbalanced (46, 47), further triggering the overwhelming production of cytokines, such as TNF-α, IL-1β, IL-6, etc. (48). The excessive inflammatory response promotes the activation of the coagulation system and the formation of microthrombosis in blood vessels, leading to disseminated intravascular coagulation (DIC) (49, 50).

“Cytokine storm” is the main cause of death in septic patients with multiple organ failure. The levels of inflammatory cytokines IL-1β, IL-6, IL-12 and IL-17 are usually high in septic patients (31, 51). In a study of community-acquired pneumonia, the mean serum concentration of IL-6 from severe septic patients was 54.1 pg/mL, compared with 36.5 pg /mL from patients who did not develop severe sepsis (52). Similarly, the serum levels of IL-1β in deceased septic patients were about 2-10 times higher than those in surviving septic patients. These clinical data suggest that high concentrations of IL-6 or IL-1β may be closely related to poor prognosis (53). Animal studies have suggested that administering anti-IL-6 antibodies reduces the expression of C5a and improves survival in septic mice (54). TNF-α is not only involved in cytokine storm but is also responsible for the induction of hypothermia and high mortality in sepsis (55). In a study of *E. coli*-induced sepsis in baboons, treatment with anti-TNF antibody at 2 h before infection protected vital organ damage and reduced mortality (56). It can be concluded from the above clinical experience that the downregulation of the excessive cytokine expression can alleviate tissue damage, reduce the conversion rate from mild to severe diseases, and improve the survival status of sepsis. In our study, DHZD could inhibit the over-activated inflammatory response (IL-6, TNF-α, IL-1β, IL-12, IFN-β and IL-10) in RAW264.7 cells, mouse primary peritoneal macrophages, and BMDMs after the stimulation of bacteria or bacterial mimic. In the early stage of sepsis, DCs can produce inflammatory factors (such as IL-6, TNF-α, IL-12 and chemokines) after bacterial phagocytosis (30). Elevated IL-12 then induces the differentiation of naive T cells into helper T cells (Th1) and the activation of NK cells, contributing to the formation of a cytokine storm (57). We demonstrated that DHZD downregulates the production of IL-6, TNF-α and IL12p70 in DCs, suggesting that it may also downregulate cytokine storm via impairing subsequent T cell over-activation.

In the LPS-induced septic mouse model, DHZD showed an impressive anti-inflammatory role and alleviated lung, liver and kidney damage, improving the survival rate. Both of the examined concentrations of DHZD showed protective effects *in vivo*. The survival rate of LPS+DHZD+DXM group was 100%, which was better than that of LPS+DXM group (60%), suggesting that the combination of DHZD and DXM may have a potential synergistic protective effect to treat sepsis.

TNF-α, interleukin, and interferonγcan induce hypothermia by regulating the thermotaxic center of the brain (58, 59). During infection, the activity of animals is reduced, which can also lead to the appearance of hypothermia (60). During sepsis, central nervous system dysfunction (88% vs 60%), elevated serum bilirubin concentration (35% vs. 15%), prolonged prothrombin time (50% vs. 23%), shock (94% vs 61%), nonrecovery from shock (66% vs. 26%), and death (62% vs. 26%), were found in patients with hypothermia compared to patients in the fever group (61). In this study, DHZD protected the loss of body temperature in CRKP-induced septic mouse model (-5.7 [vs -2.6 [) compared to those of CRKP group mice, suggesting that DHZD-treated infective mice have a better prognosis.

The pathogen load is usually related to the severity of infectious diseases. Macrophage is a ubiquitous phagocytic cell type of innate immune system involved in defensive inflammatory response and pathogen clearance during bacterial infection (62, 63). Besides macrophages, neutrophils and DCs are also important phagocytic cells in the body, which can phagocytize bacteria and decrease the bacterial load. Our study used macrophages as a cell model to investigate whether DHZD can influence cellular phagocytosis. DHZD promoted phagocytosis of bacteria in a clathrin-and-actin-dependent manner.

CD36, a class B scavenger receptor family member, is expressed in monocytes, macrophages, endothelial cells, and adipocytes (52). The upregulation of CD36 expression by DHZD might at least partially explain the enhanced phagocytosis of macrophages. Not only the expression of phagocytic receptors of phagocytes but also the proportions and numbers of neutrophils and monocytes/macrophages in lungs were increased by DHZD after the LPS challenge, suggesting that DHZD might contribute to a quick control of bacterial load at an early stage of infection through phagocytosis. Accordingly, DHZD might be protective during bacterial infection and decrease organ damage.

As a member of innate immune cells, DCs have an important role in the line of defense against pathogens. Immature DCs have a strong phagocytic ability. As they mature, the phagocytic ability of DC is decreased, and the expression of MHC II molecules and co-stimulatory molecules (CD86, CD80, and CD40) are increased, accompanied by the increased antigen presenting ability to evoke T cell activation (29, 64, 65). In our study, the downregulated expression of co-stimulatory molecules (CD80, CD86 and CD40) and MHC II by DHZD kept DCs in an immature status, helping to maintain their strong phagocytic ability and prevent the over-activation of T cells.

The PI3K/Akt signaling pathway is an important regulator of cytokine production after TLR4 ligation with LPS during G^-^ bacterial infection (66). Previous studies have shown that the inhibition of PI3K/Akt activation in macrophages or DCs reduces the secretion of cytokines (IL-6, TNF-α, IL-12, IL-10 and IL-1β) (42, 67). Though IL-10 has a well-known role in anti-inflammation, extremely high levels of IL-10 are harmful to patients. For example, previous studies reported 3.3 times higher serum IL-10 levels in septic patients who died compared to survivors (68). In this study, DHZD inhibited the activation of the PI3K/Akt/p70S6K signaling pathway but did not influence the activation of ERK and p38 MAPK signaling pathways, accounting for its anti-inflammatory role.

In conclusion, DHZD exerts an obvious anti-inflammatory effect *in vitro* and *in vivo* by targeting the host immune response and thus may be a potential therapeutic drug for treating sepsis caused by bacterial infection and even drug-resistant bacterial infection. Further experiments are needed to assess whether the phagocytic ability of bacteria besides *E. coli* could also be increased by DHZD. Interestingly, the combination of DHZD and DXM is expected to improve the survival rate of septic mice, hinting that DHZD might be a potential adjuvant drug candidate for the treatment of sepsis.

## Acknowledgments

This work was supported by grants from National Key Research and Development Program of China (2021YFE0200900), National Natural Science Foundation of China (82073901, 82073726), Shanghai Municipal Science and Technology Major Project (ZD2021CY001), the Key Research and Development Program of Shaanxi Province (2020SF-226), three-year Action Plan for Shanghai TCM Development and Inheritance Program [ZY (2021–2023)-0401 and ZY (2021–2023)-0103]. Traditional Chinese Medicine Research Project of Shanghai Municipal Health Commission (2022QN043). We are thankful for Dr. Yi-jian Chen from Huashan Hospital affiliated to Fudan University to provide Carbapenem-resistant *Klebsiella pneumoniae* (CRKP, HS11286). Thank Ali Mollahassani for language polishing of this paper.

## Author contribution

Yue-juan Zheng designed the experiments. Xiang-yang Qin modified and provided DHZD. Ying-xiang Zou, Tian-nan Xiang, Huan Zhang, Yu-he Ma, Xiao Wu, Qi-lin Huang, Biao Lei and Jing-wen Mu performed the laboratory assays. Ying-xiang Zou, Tian-nan Xiang, Huan Zhang and Yu-he Ma analyzed the data. Yue-juan Zheng and Ying-xiang Zou wrote the manuscript. Tian-nan Xiang and Xiang-yang Qin contributed to manuscript revision. All authors read and approved the manuscript.

