## supplemental material for "Dehydrozaluzanin C- derivative protects septic mice by alleviating over-activated inflammatory response and promoting the phagocytosis of macrophages"

**Supplementary information**

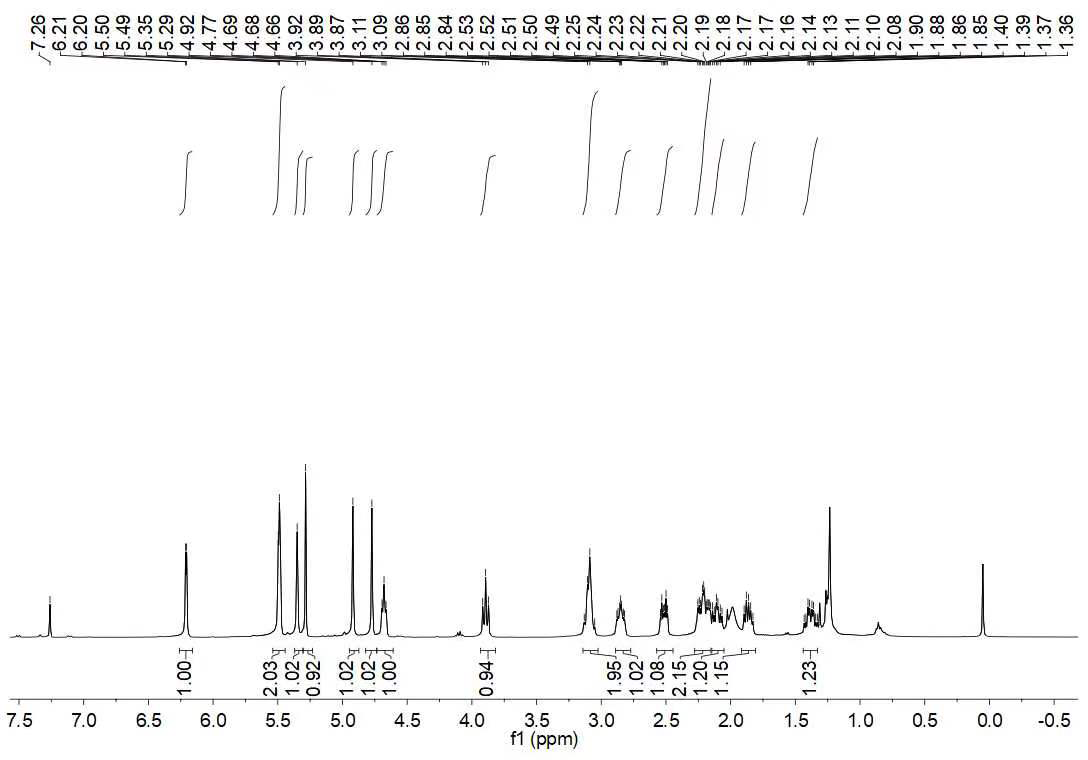

**Supplementary Figure 1.** **Plot of the spectral data of DHZD.**

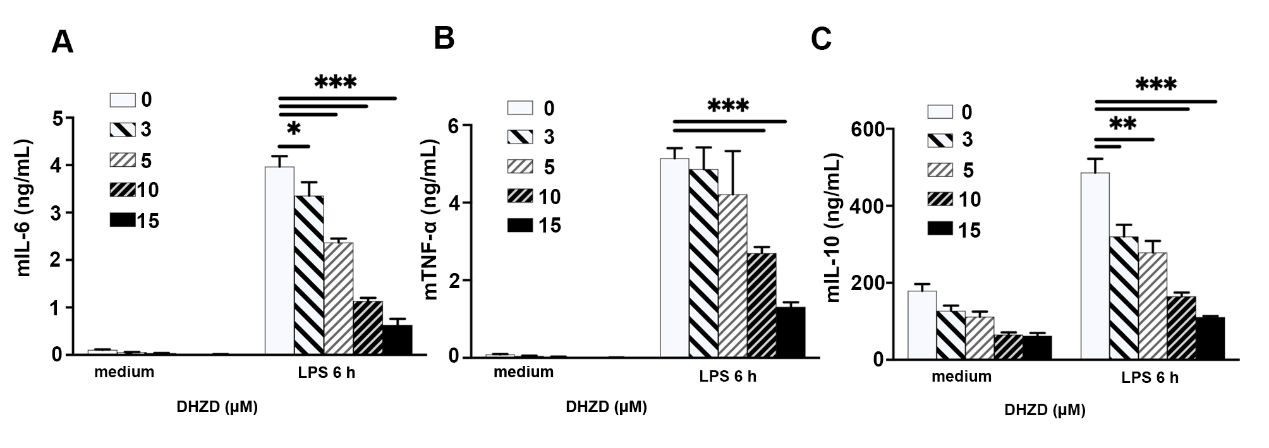

**Supplementary Figure 2. DHZD decreases the secretion of LPS- induced IL-6, TNF-α, MCP-1 and IL-10 in BMDMs.** The secretion of IL-6 (A), TNF-α (B) and IL-10 (C) were detected by ELISA. Data were shown as mean ± SD from three independent experiments. *, *P* <0.05, **, *P* <0.01, and ***, *P* <0.001.

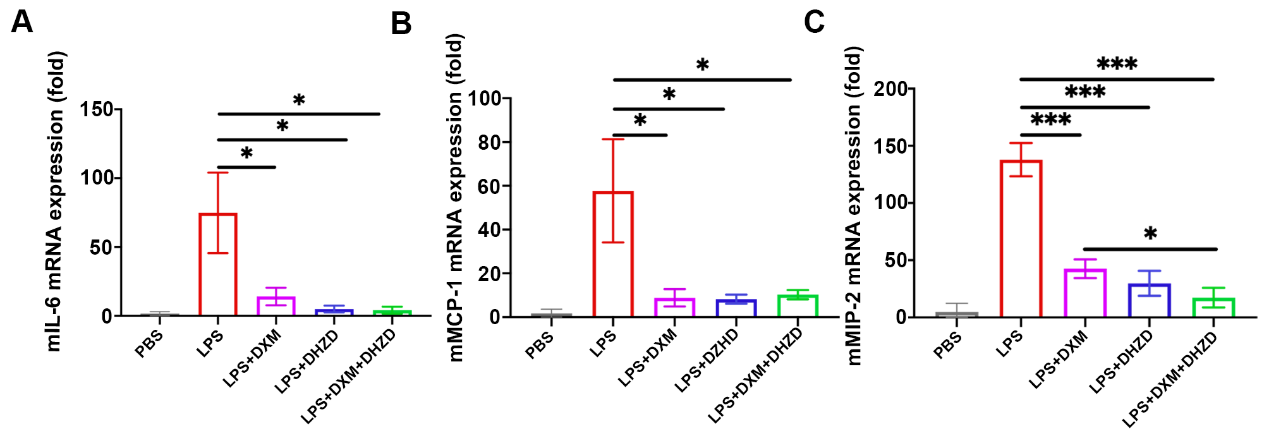

**Supplementary Figure 3. DHZD deceases the expression of inflammatory response in lung of LPS-induced acute peritonitis mice.** The mRNA expression of IL-6 (A), MCP-1(B) and MIP-2 (C) in lung tissues were detected by qRT-PCR. Data were shown as mean ± SD of three mice per group. *, *P* <0.05, and ***, *P* <0.001.
